# Genome comparisons indicate recent transfer of *w*Ri-like *Wolbachia* between sister species *Drosophila suzukii* and *D. subpulchrella*

**DOI:** 10.1101/135475

**Authors:** William R. Conner, Mark L. Blaxter, Gianfranco Anfora, Lino Ometto, Omar Rota-Stabelli, Michael Turelli

**Affiliations:** Department of Evolution and Ecology, University of California, One Shields Avenue, Davis, California 95616, USA; Institute of Evolutionary Biology and Edinburgh Genomics Facility, University of Edinburgh, Edinburgh EH9 3JT, UK; Chemical Ecology Lab, Department of Sustainable Agro-ecosystems and Bio-resources, Fondazione Edmund Mach, San Michele all’Adige (TN), Italy; Centre Agriculture Food Environment, University of Trento, San Michele all’Adige (TN), Italy

**Keywords:** spotted-wing Drosophila, introgression, horizontal transmission, molecular clocks, relative rates, cytoplasmic incompatibility loci

## Abstract

*Wolbachia* endosymbionts may be acquired by horizontal transfer, by introgression through hybridization between closely related species, or by cladogenic retention during speciation. All three modes of acquisition have been demonstrated, but their relative frequency is largely unknown. *Drosophila suzukii* and its sister species *D. subpulchrella* harbor *Wolbachia*, denoted *w*Suz and *w*Spc, very closely related to *w*Ri, identified in California populations of *D. simulans.* However, these variants differ in their induced phenotypes: *w*Ri causes significant cytoplasmic incompatibility (CI) in *D. simulans*, but CI has not been detected in *D. suzukii* or *D. subpulchrella.* Our draft genomes of *w*Suz and *w*Spc contain full-length copies of 703 of the 734 single-copy genes found in *w*Ri. Over these coding sequences, *w*Suz and *w*Spc differ by only 0.004% (i.e., 28 of 704,883 bp); they are sisters relative to *w*Ri, from which each differs by 0.014-0.015%. Using published data from *D. melanogaster, Nasonia* wasps and *Nomada* bees to calibrate relative rates of *Wolbachia* versus host nuclear divergence, we conclude that *w*Suz and *w*Spc are too similar - by at least a factor of 100 - to be plausible candidates for cladogenic transmission. These three *w*Ri-like *Wolbachia*, which differ in CI phenotype in their native hosts, have different numbers of orthologs of genes postulated to contribute to CI; and the CI loci differ at several nucleotides that may account for the CI difference. We discuss the general problem of distinguishing alternative modes of *Wolbachia* acquisition, focusing on the difficulties posed by limited knowledge of variation in absolute and relative rates of molecular evolution for host nuclear genomes, mitochondria and *Wolbachia*.

## Introduction

*Drosophila suzukii* Matsumura (Diptera Drosophilidae) is an invasive and destructive fruit fly native to South East Asia that has recently invaded North America, South America and Europe (Hauser 2011; Cini *et al.* 2012; Rota-Stabelli *et al.* 2013). While most *Drosophila* species oviposit in fermenting fruits, *D. suzukii* and its close relative *D. subpulchrella* Takamori and Watabe use their atypical serrated ovipositors to pierce the skin of ripening soft fruits and lay eggs in them (Fig. 1, McEvey 2017a,.b; Atallah *et al.* 2014). Leveraging the genetic resources of *D. melanogaster, D. suzukii* and *D. subpulchrella* (both members of the *D. melanogaster* species group) are becoming model species for fundamental and applied studies.

**FIGURE 1.**
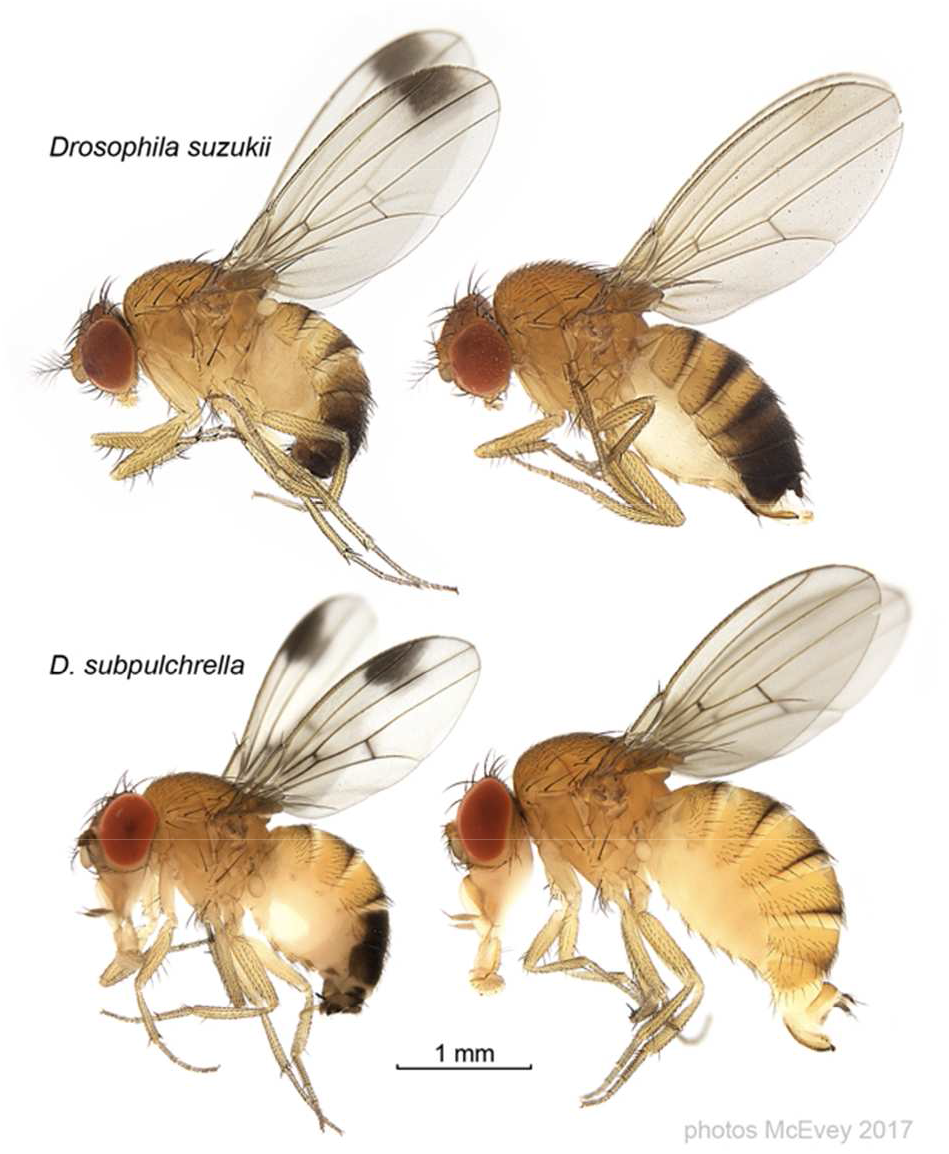
*Drosophila suzukii* and *D. subpulchrella*, with males on the left. The photos are from McEvey (2017a,b), the composite image is courtesy of Shane McEvey.

*Wolbachia* are obligately intracellular, maternally inherited alpha-proteobacteria found in about half of all insect species and many other terrestrial arthropods and nematodes (Weinert *et al.* 2015). *Wolbachia* are often associated with reproductive manipulations, including cytoplasmic incompatibility (CI) (Hoffmann & Turelli 1997), male killing (Hurst & Jiggins 2000), feminization (Rousset *et al.* 1992) and parthenogenesis induction (Stouthamer *et al.* 1993), all of which enhance the relative fitness of infected females. But many *Wolbachia* infections, including those in *D. suzukii* and its sister species *D. subpulchrella*, cause no detectable reproductive manipulation and presumably persist by enhancing host fitness (Kriesner *et al.* 2013; Hamm *et al.* 2014; Mazzetto *et al.* 2015; Kriesner *et al.* 2016; Cattel *et al.* 2016). Indeed, it seems increasingly plausible that even infections that cause reproductive manipulations become established in new hosts because they enhance fitness, and hence tend to increase in frequency even when very rare (Kriesner *et al.* 2013). For example, the most common *Wolbachia* reproductive manipulation is CI, in which embryos produced by uninfected females mated with infected males suffer increased mortality. Because CI is essentially irrelevant to the frequency dynamics of rare infections, initial spread of both CI-causing infections and infections that do not manipulate reproduction is likely to be driven by mutualistic effects such as fecundity enhancement (Weeks *et al.* 2007; Fast *et al.* 2011), protection from viruses (Teixeira *et al.* 2008) or metabolic provisioning (Brownlie *et al.* 2009).

To understand why *Wolbachia* are found in so many species, it is critical to know how *Wolbachia* infections are acquired and how long *Wolbachia-host* associations persist. As noted by Raychoudhury *et al.* (2008), although *Wolbachia* are maternally transmitted, host lineages can acquire *Wolbachia* in three ways: by cladogenic transmission, in which an infection persists through speciation; by introgression, in which hybridization of closely related species leads to maternal cytoplasm transfer; or by horizontal transmission, in ways that remain indeterminate, in which *Wolbachia* are transferred between closely or distantly related species through non-sexual mechanisms (such as predation or parasitism).

To complement an analysis of *Wolbachia* population biology and effects in *Drosophila suzukii* and its sister species *D. subpulchrella*, Hamm *et al.* (2014) presented a meta-analysis of *Wolbachia* infections in *Drosophila* species that addressed the frequency of both reproductive manipulation and alternative modes of acquisition. However, we show that their informal methodology underestimated the relative frequencies of horizontal and introgressive transmission. Horizontal transmission of *Wolbachia* was first demonstrated by extreme discordance of the phylogenies of distantly related hosts and their infecting *Wolbachia* (O’Neill *et al*. 1992). In contrast, horizontal transmission seems negligible within the two species that have been examined most intensively, *D. simulans* (Turelli & Hoffmann 1995) and *D. melanogaster* (Richardson *et al.* 2012). Hamm *et al.* (2014) implicitly assumed that if two closely related host species share closely related *Wolbachia*, the infections are likely to have been acquired by either cladogenic transmission or introgression. In particular, Hamm *et al.* (2014) postulated that because *D. suzukii* and its sister *D. subpulchrella* have concordant mitochondrial and nuclear phylogenies and harbor very similar *Wolbachia*, as indicated by identity at the Multi Locus Sequence Typing (MLST) loci used to classify *Wolbachia* (Baldo *et al.* 2006), cladogenic *Wolbachia* acquisition was likely. Here we use comparative analyses of draft *Wolbachia* genomes, and extensive nuclear data from *Drosophila* and other insect hosts, to refute this hypothesis.

The three alternative modes of *Wolbachia* acquisition would be trivial to distinguish if reliable chronograms (dated phylogenies) were available for the nuclear, mitochondrial and *Wolbachia* genomes. Under cladogenic transmission, without subsequent introgression or horizontal transmission, roughly concordant chronograms for all three genomes are expected. From the arguments of Gillespie & Langley (1979), we expect a slightly longer divergence time for nuclear than mitochondrial or *Wolbachia* given the greater intraspecific variation observed in nuclear DNA. However, for typical pairs of *Drosophila* species that diverged on the order of 10^6^ years ago (Coyne & Orr 1989, 1997), this discordance under cladogenic acquisition is unlikely to be as large as a factor of two. Under introgression without subsequent horizontal transmission, the mitochondrial and *Wolbachia* chronograms should be concordant (because they are simultaneously maternally transmitted) and show more recent divergence than the bulk of the nuclear genome. Finally, under horizontal transmission, more recent divergence is expected between infecting *Wolbachia* than between either the host mitochondrial or nuclear genomes. These simple criteria are difficult to apply because of uncertainty concerning the relative rates of nuclear, mitochondrial and *Wolbachia* divergence. Here, using all available comparative data for *Wolbachia* and host divergence, we conclude that the *Wolbachia* in *D. suzukii* and *D. subpulchrella* are far too similar to make cladogenic transmission plausible. Our conclusion does not exclude the possibility that *D. suzukii* and *D. subpulchrella* retained a *Wolbachia* infection from their common ancestor. Our data indicate only that their current *Wolbachia* are too similar to have been diverging since the speciation of their hosts. In principle, one could establish cladogenic transmission followed by introgression or horizontal transmission if traces of historical infections could be found in host genomes (Hotopp *et al.* 2007). Unfortunately, as shown below, no such traces were found in our *D. suzukii* or *D. subpulchrella* genomes.

In addition to assessing *Wolbachia* acquisition, we examine patterns of molecular evolution by comparing the draft genomes for *w*Suz (Siozos *et al.* 2013) and *w*Spc (this paper) to the *w*Ri reference genome (Klasson *et al.* 2009). We consider both a general pattern, namely the relative frequencies of non-synonymous and synonymous substitutions, and sequence divergence for candidate loci associated with two *Wolbachia*-induced phenotypes, life shortening and CI. The “Octomom” duplication, which distinguishes *w*MelPop (Min & Benzer 1997) from *w*Mel (Wu *et al.* 2004), contains the genes *WD0507-WD0514* and is associated with extremely high *Wolbachia* titer and life shortening in *D. melanogaster* (Chrostek & Teixeira 2015, but see Rohrscheib *et al.* 2016 for a critique and LePage *et al.* 2017 and Chrostek & Teixeira 2017 for support of the hypothesis connecting these loci to life shortening or *Wolbachia* titer). Beckmann & Fallon (2013) used proteomics to identify the locus wPip_0282 in *w*Pip, the *Wolbachia* found in *Culex pipiens*, as a candidate for producing CI. They found at least one homolog of this WO prophage locus in several CI-causing *Wolbachia*, including *w*Mel and *w*Ri. Within *w*Pip and other *Wolbachia* genomes, *w*Pip_0282 and each homolog seemed to be part of two-gene operons, with **w*Pip_0282* adjacent to **w*Pip_0283.* This pair is orthologous to *WD0631* and *WD0632* in *w*Mel, and there are three homologous/paralogous pairs in *w*Ri. Beckmann *et al.* (2017) and LePage *et al.* (2017) provide experimental and bioinformatic evidence that *WD0631* and *WD0632* contribute to CI (but LePage *et al.* [2017] argue against the operon hypothesis). We examine differences in homologs and paralogs of these loci among *w*Suz, *w*Spc and *w*Ri.

## Materials and methods

### Sequence data

Genome data for *D. suzukii* and *D. subpulchrella* were generated by Edinburgh Genomics. The *D. suzukii* genome data were generated from an inbred Italian line (the Trento strain) as presented in Ometto *et al.* (2013), with the *Wolbachia*, *w*Suz, presented in Siozos *et al.* (2013). Illumina HiSeq2000 120-base, paired-end sequence data were generated from two libraries of 180 and 300 base pair (bp) inserts. The *D. subpulchrella* genome data were generated from a stock maintained at the Fondazione Edmund Mach lab that was established from San Diego Drosophila Species Stock center strain 14023-0401.00, originally from Japan. Illumina HiSeq2000 125-base, paired-end sequence data were generated from two libraries of 350 and 550 bp inserts.

### *Assembly of* Wolbachia *in* D. subpulchrella

To assemble *w*Spc, we initially cleaned, trimmed and assembled reads for the *Wolbachia-* infected *D. subpulchrella* using Sickle (https://github.com/najoshi/sickle) and SOAPdenovo v. 2.04 (Luo *et al.* 2012). For the assembly, *K* values of 31, 41, … and 101 were tried, and the best assembly (fewest contigs and largest N50) was kept. This preliminary assembly had over 100,000 contigs with a total length of 243 megabases (Mbp). Details of the *D. subpulchrella* assembly will be published elsewhere, together with a comparison to the *D. suzukii* genome. Most of the contigs were identified through BLAST search as deriving from *Drosophila.* Minor contamination from microbiota (such as *Acetobacter* spp.) was identified. Contigs with best nucleotide BLAST matches (with E-values less than 10^−10^) to known *Wolbachia* sequences were extracted as the draft assembly for *w*Spc. We also attempted filtering the reads by alignment to *w*Ri and assembling with SPAdes 3.0 (Bankevich *et al.* 2012). The assembly of *w*Spc is available from GenBank under accession number XXXXX [to be advised].

To assess the quality of our draft *w*Spc and *w*Suz assemblies, we used BUSCO v. 3.0.0 (Simão *et al.* 2015) to search for orthologs of the near-universal, single-copy genes in the BUSCO proteobacteria database. As a control, we performed the same search using the complete reference genomes for *w*Ri (Klasson *et al.* 2009), *w*Au (Sutton *et al.* 2014), *w*Mel (Wu *et al.* 2009), *w*Ha and *w*No (Ellegaard *et al.* 2013).

### Phylogeny and estimates of divergence of wSpc and wSuz

The *Wolbachia* MLST loci *gatB, hcpA, coxA, fbpA, and ftsZ* (Baldo *et al.* 2006) were identified in the assemblies using BLAST. As reported in Hamm *et al.* (2014), the MLST sequences from *w*Spc and *w*Suz were identical both to each other and to those of the *w*Ri reference genome from *D. simulans* (Klasson *et al.* 2009).

To distinguish these *Wolbachia* and determine their relationships, we extracted additional orthologous loci from the draft genomes. We annotated the genomes of *w*Suz and *w*Spc with Prokka v 1.11 (Seemann 2014), which identifies orthologs to reference bacterial genes. To normalize our comparisons, we also annotated the genomes of *w*Ri (Klasson *et al.* 2009), *w*Au (Sutton *et al.* 2014) and *w*Mel (Wu *et al.* 2004; Richardson *et al.* 2012). We selected 512 genes present in full length and single copy in all five genomes, avoiding incomplete or pseudogenes and loci with paralogs. Genes were treated as single copy if no other gene in the genome was matched to the same reference bacterial gene by Prokka, and as full length if the orthologs in the other *Wolbachia* genomes all had the same length. The nucleotide sequences of the genes were aligned with MAFFT v. 7 (Katoh 2013) and concatenated, giving an alignment of 480,831 bp. The strain phylogeny was estimated with a phylogram constructed with MrBayes v. 3.2 (Ronquist & Huelsenbeck 2003) using the GTR+Γ model, partitioned by codon position. All model parameters for each partition were allowed to vary independently, except topology and branch length. We ran two independent chains, each with four incrementally heated subchains, for 1,000,000 generations. Trace files for each analysis were visualized in Tracer v. 1.6 (Rambaut *et al.* 2014) to ensure convergence of all continuous parameters. The first 25% of the generations were discarded as burn-in. Only one topology had posterior probability > 0.001.

To estimate the divergence between *w*Suz and *w*Spc, 703 genes present in full length and single copy in *w*Suz, *w*Spc, and *w*Ri (spanning a total of 704,883 bp) were extracted and aligned with MAFFT v. 7. As an additional assessment of the completeness of the *w*Suz and *w*Spc assemblies, we calculated the number of single-copy genes in the *w*Ri reference and found 734. The resulting alignments were concatenated. To estimate a chronogram, we assumed for simplicity that each partition evolved at a constant rate across the tree (allowing the rates to differ among codon positions). The constant-rate chronogram was estimated with MrBayes v. 3.2, using the same procedure as our five-sequence *Wolbachia* phylogram (which included *w*Mel and *w*Au). The age of the *w*Suz-*w*Spc node was set at 1, as an arbitrary scaling of relative ages.

### *Nuclear divergence between* D. subpulchrella *and* D. suzukii

Hamm *et al.* (2014) used *Drosophila* nuclear data extracted from Yang *et al.* (2012) to assess the relationships of *D. suzukii, D. subpulchrella* and *D. biarmipes*, but these data have subsequently been shown to be unreliable (Catullo & Oakeshott 2014). We reassessed these relationships and compared the *Wolbachia* and nuclear chronograms for *D. suzukii* and *D. subpulchrella.* We identified in FlyBase complete coding regions for *D. melanogaster* for the ten nuclear loci used by Hamm *et al.* (2014) *(H2A, Adh, amylase, amyrel, cdc6, ddc, esc, hb, nucl*, and ptc), plus ten additional nuclear loci *(aconitase, enolase, glyp, glys, pepck, pgi, pgm, tpi, white*, and wg). We used BLAST to identify orthologs in the *D. suzukii* assembly of Ometto *et al.* (2013), the unpublished draft *D. subpulchrella* assembly described above, a *D. biarmipes* assembly (Chen *et al.* 2014), and a second-generation *D. simulans* assembly (Hu *et al.* 2012). Data for *H2A* and *amylase* were eliminated because *H2A* had multiple non-identical paralogs in each species and homologs of *D. melanogaster amylase* could not be found in the assemblies. The coding sequences for the remaining 18 loci were aligned with MAFFT v. 7 and concatenated. (Our nuclear data from *D. subpulchrella* are available from GenBank under accession numbers MF908506-MF909523.) The alignment was analyzed with MrBayes v. 3.2 using the same model and procedures used for our *Wolbachia* analyses, except that we partitioned the data by both gene and codon position. We estimated both a phylogram and a constant-rate chronogram. The latter assumed that each partition evolved at a constant rate over the tree. The age of the most recent common ancestor (MRCA) of *D. suzukii* and *D. subpulchrella* was set at 1, as an arbitrary scaling of relative ages. To test the robustness of our relative divergence-time estimates for the host species, we also estimated the chronogram using a relaxed-clock model in RevBayes (Hoehna *et al.* 2016). This analysis also partitioned the data by gene and codon position and used the GTR+Γ model, but it assumed uncorrelated lognormal rate variation across branches. Following the RevBayes tutorial (https://github.com/revbayes/revbayes_tutorial/blob/master/RB_BayesFactor_Tutorial/scripts/marginal_likelihood_GTR_Gamma_inv.Rev), we used a lognormal prior with mean and standard-deviation parameters (-*X*^2^/2, *X*) and a lognormal hyperprior on *X* with parameters (ln(2)/4, Sqrt[ln(2)/2]). To estimate *k*_s_ and *k*_a_ between *D. suzukii* and *D. subpulchrella*, we used DNAsp v. 5.10 (Rozas 2009).

Following Hotopp *et al.* (2007), we looked for evidence of genetic transfer from *w*Suz and *w*Spc (or other *Wolbachia)* to these hosts. The *D. suzukii* and *D. subpulchrella* assemblies (including the *Wolbachia* contigs) were BLASTed against both all known *melanogaster* group nuclear sequences and all known *Wolbachia* sequences. We sought contigs for which part mapped to a *Drosophila* nuclear sequence and not to any *Wolbachia* sequence while another part mapped to a *Wolbachia* sequence and not to any *Drosophila* nuclear sequence.

### Analysis of divergence between wSpc, wSuz and wRi

The trimmed Illumina reads from *D. suzukii* and *D. subpulchrella* were aligned to the *w*Ri reference (Klasson *et al.* 2009) with bwa v. 0.7.12 (Li & Durbin 2009). As a control, we also aligned Illumina reads from Riv84 (Iturbe-Ormaetxe *et al.* 2010), the *D. simulans* line used to make the *w*Ri reference. Normalized read-depth for each alignment was calculated over sliding 1000 bp windows by dividing the average depth in the window by the average depth over the entire genome. Putative copy number variant (CNV) locations were identified with ControlFREEC v. 8.0 (Boeva *et al.* 2012), using 500 bp windows and the Riv84 alignment as a control. For the bulk of the genomes, we used an expected ploidy of one, but for variants involving sequences duplicated in *w*Ri, we used a ploidy of two. We calculated *P*-values for each putative CNV using the Kolmogorov-Smirnov test implemented in ControlFREEC.

Sequences for the “Octomom” genes *WD0507-WD0514* (Chrostek & Teixeira 2015, *cf.* Rohrscheib *et al.* 2016; Chrostek & Teixeira 2017) were extracted from the *w*Mel reference (Wu *et al.* 2004; Richardson *et al.* 2012). Using BLAST, we identified orthologs in the *w*Ri reference (Klasson *et al.* 2009) and the draft assemblies for *w*Suz and *w*Spc.

Sequences homologous to loci putatively involved in CI in other *Wolbachia* strains (Beckmann & Fallon 2013; LePage *et al.* 2017; Beckmann *et al.* 2017) were extracted from *w*Ri (Klasson *et al.* 2009) and the draft assemblies for *w*Suz and *w*Spc. Differences among these three genomes at these loci were assessed by aligning the *w*Suz and *w*Spc reads to the *w*Ri reference and calculating the percentage of reads with the non-*w*Ri base.

To identify a specific insertion of the transposable element ISWpi7, which occurs in 21 identical copies in *w*Ri, and whose position differentiates *w*Spc and *w*Suz from *w*Ri, an additional assembly step was required. The novel insertion occurs in the *w*Spc and *w*Suz orthologs of *WRi_006720*, one of the CI-associated loci discussed below. The *D. suzukii* and *D. subpulchrella* reads were aligned to the *w*Spc assembly with bwa 0.7.12 (Li & Durbin 2009). For both contigs that contain part of the *WRi_006720* gene, reads mapping to the ISWpi7 transposable element plus the neighboring 500 bp were extracted and assembled with SOAPdenovo v. 2.04 (Luo *et al.* 2012), using a *K* value of 55. Both the *D. suzukii* and *D. subpulchrella* reads assembled into a single contig containing the two pieces of *WRi_006720* interrupted by a single copy of ISWpi7. To test this bioinformatic result, we designed two pairs of PCR primers that spanned the hypothesized junctions between the ortholog of *WRi_006720* and ISWpi7. For the first set of primers (forward: ATGGTCACATTGAACAGAGGAT, reverse: GTTGGTGCTGCAATGCGTAA), the forward primer attaches at 728945-728966, part of *WRi_006720.* For the second set of primers (forward: AGCGTTGTGGAGGAACTCAG, reverse: CGTCATGCTGCAGTGCTTAG), the reverse primer attaches at 729570-729589, part of *WRi_006720.* No detectable product is expected with either primer set in *w*Ri, which does not contain the insert in *WRi_006720*; whereas each primer set is expected to produce a unique band with *w*Spc and *w*Suz.

## Results

### *Draft genome assembly for* wSpc, *the* Wolbachia *from* D. subpulchrella

We generated a draft assembly of *w*Spc by filtering contigs from a joint *Wolbachia-D. subpulchrella* assembly. The draft *w*Spc assembly was in 100 contigs with N50 length of 31,871 bp and total length of 1.42 Mbp. This length is close to the 1.45 Mbp *w*Ri reference (Klasson *et al*. 2009), suggesting that it may represent a nearly complete genome. In contrast, the assembly produced by SPAdes 3.0 had N50 of 8,360 bp and total length of 1.20 Mbp.

Out of 221 near-universal, single-copy orthologs in proteobacteria, BUSCO 3.0.0 (Simão *et al.* 2015) found effectively the same number in all of the tested genomes (*w*Ri, *w*Au, *w*Mel, *w*Ha, *w*No, and the drafts of *w*Suz and *w*Spc). Our draft assemblies for *w*Spc and *w*Suz contain two BUSCO-annotated genes not found in *w*Ri and *w*Mel. See Table S1 for detailed information.

### Wolbachia divergence

We aligned and compared *w*Spc and *w*Suz at 703 protein-coding loci (704,883 bp) and identified only 28 single-nucleotide variants (SNV), an overall divergence of 0.004%. *w*Suz had 103 SNV compared to *w*Ri (0.015% divergence) and *w*Spc had 99 SNV (0.014% divergence) (Table S2). Most (87) of these SNV are shared. There were too few differences to definitively determine whether these genomes are recombinant (Ellegaard *et al.* 2013), but the data were fully consistent with no recombination (*i.e*., with so few differences, we have no power to detect recombination). Bayesian phylogenetic analysis placed *w*Suz and *w*Spc as sisters relative to *w*Ri (Fig. 2A). For *w*Suz and *w*Spc, we derived point estimates and 95% confidence intervals for divergence at each codon position, calculated as the rate multiplier for that position times the branch length (fixed to 1) (Table 1). The rate multipliers express the relative rate of evolution for each codon position. Hence, the expected number of substitutions for each codon position along each branch of the phylogram is the branch length times the rate multiplier for that position. The estimated chronogram (Fig. 2B) shows that the divergence time of *w*Ri from its MRCA with *w*Spc and *w*Suz is 3.51 times the divergence time of *w*Spc and *w*Suz, with a 95% confidence interval of (2.41, 4.87). We found no difference in the rates of divergence for first, second and third codon positions, as also observed in the co-divergence of *Wolbachia* and mtDNA haplotypes in *D. melanogaster* (Richardson *et al.* 2012). Following from this, estimates of synonymous, *k*_s_, and non-synonymous, *k*_a_, substitution rates were very similar (Table 1).

**FIGURE 2.**
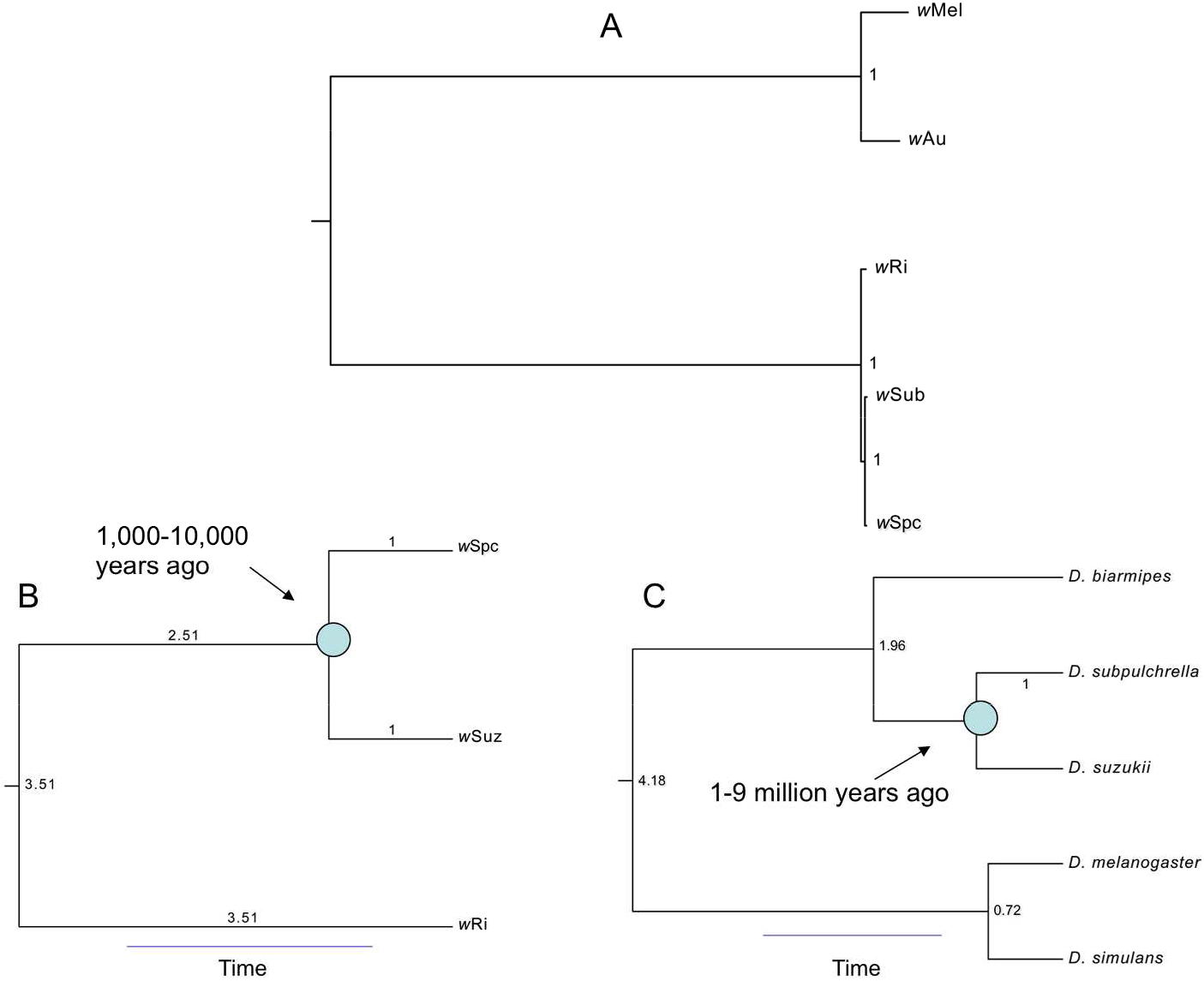
Phylogram and chronograms for the *Wolbachia* and hosts discussed. Clade posterior probabilities are shown. A) *Wolbachia* phylogram. B) *Wolbachia* chronogram with an estimate of the divergence time for *w*Suz and *w*Spc. Branch lengths relative to the *w*Spc-*w*Suz divergence are shown. All clade posterior probabilities are 1.0. C) Host chronogram with an estimate of divergence time for *D. suzukii* and *D. subpulchrella.* Branch lengths relative to the *D. suzukii-D. subpulchrella* divergence are shown. All clade posterior probabilities are 1.0.

**Table 1.** Estimated number of substitutions per site by codon position between *w*Suz and *w*Spc, plus estimates of synonymous (*k*_s_) and non-synonymous (*k*a) substitution rates, see the text for details.

| Position | Point Estimates | 95% Confidence Interval |
| --- | --- | --- |
| 1<sup>st</sup> | $5.0\times10^{-5}$ | $(3.0\times10^{-5}, 7.0\times10^{-5})$ |
| 2<sup>nd</sup> | $3.2\times10^{-5}$ | $(1.6\times10^{-5}, 4.6\times10^{-5})$ |
| 3<sup>rd</sup> | $4.0\times10^{-5}$ | $(2.4\times10^{-5}, 5.6\times10^{-5})$ |
| Overall ($k_s, k_a$) | $(3\times10^{-5}, 4\times10^{-5})$ | |

### Host divergence

The host phylogram (not shown) and chronogram (Fig. 2C) demonstrate that *D. subpulchrella* and *D. suzukii* are sisters relative to *D. biarmipes*, as reported by Hamm *et al.* (2014). The divergence time of *D. biarmipes* from its MRCA with *D. subpulchrella* and *D. suzukii* was estimated to be 1.96 times the divergence time for *D. subpulchrella* and *D. suzukii*, with 95% confidence interval (1.84, 2.08). The *D. melanogaster* and *D. simulans* divergence time estimate is 0.72 times as large as the estimated divergence time for *D. subpulchrella*-*D. suzukii*, with a 95% confidence interval of (0.65, 0.78). Point estimates and 95% confidence intervals for divergence at each codon position between *D. subpulchrella* and *D. suzukii* were calculated as the rate multiplier for that position times the branch length (fixed to 1) (Table 2). Our estimate of the third-codon-position substitutions per site, which we use to date *D. subpulchrella-D. suzukii* divergence, is 9.20×10^−2^ and a 95% confidence interval of (8.6×10^−2^, 9.80×10^−2^).

**Table 2.**
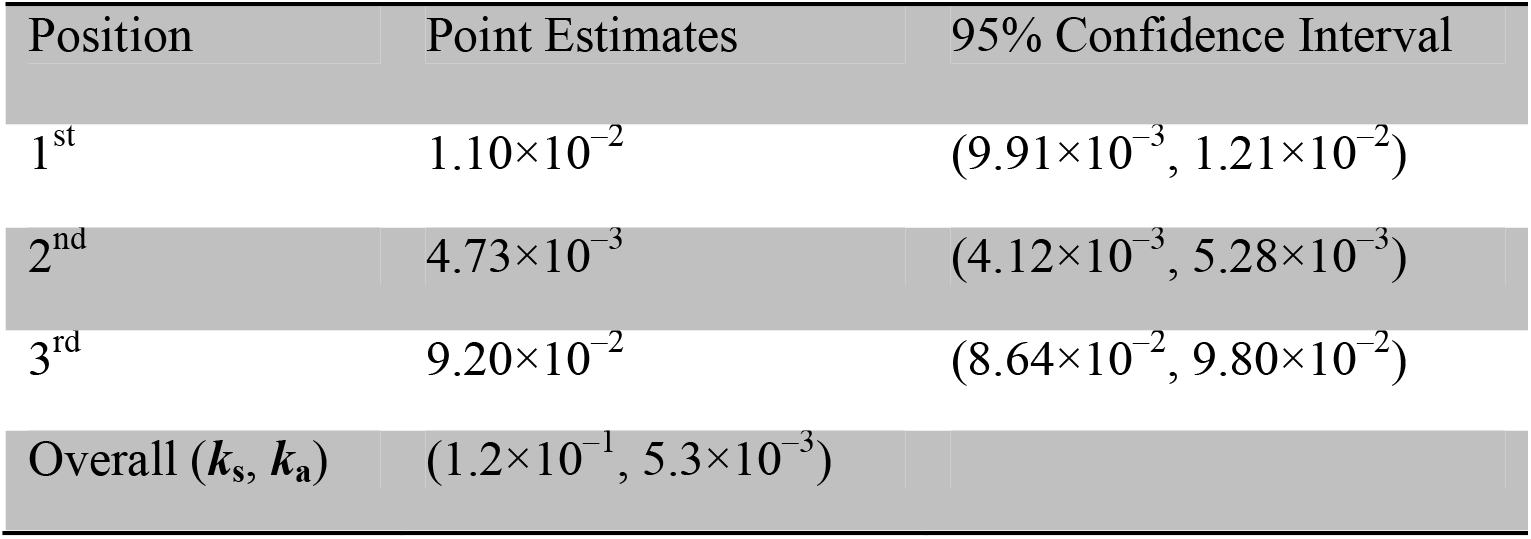
Estimated number of substitutions per site by codon position between *D. suzukii* and *D. subpulchrella* for 18 nuclear loci, plus estimates of synonymous (*k*_s_) and non-synonymous (*k*_a_) substitution rates, see the text for details.

Our RevBayes (2016) relaxed-clock chronogram (not shown) was very similar to Fig 2C. The divergence time of *D. biarmipes* from its MRCA with *D. subpulchrella* and *D. suzukii* was estimated to be 1.84 times the divergence time for *D. subpulchrella* and *D. suzukii* (instead of 1.96). Similarly, the *D. melanogaster* and *D. simulans* divergence time estimate is 0.76 times as large as the estimated divergence time for *D. subpulchrella-D. suzukii* (instead of 0.72). We note that the model underlying this analysis assumes for computational convenience that each partition undergoes proportional rate variation across each branch, *i.e.*, each partition speeds up or slows down by the same amount along each branch (but see Langley & Fitch 1974).

We found no evidence for partial integration of any *Wolbachia* sequence into the nuclear genomes of either *D. subpulchrella* or *D. suzukii*.

### Calibrations for Wolbachia versus host genome divergence and interpretation

We used estimates of relative divergence of the *Wolbachia* and *Drosophila* genomes to assess cladogenic versus lateral transmission of *w*Spc and *w*Suz. Our strategy was to compare our estimates of relative *Wolbachia/host* divergence to ratios obtained from published examples of cladogenic *Wolbachia* transmission. Table 3 summarizes our data and the data from two *Nasonia* wasp species (Raychoudhury *et al.* 2008, *w*NlonB1 versus *w*NgirB) and four *Nomada* bee species (Gerth & Bleidorn 2016, plus unpublished data kindly provided by the authors). Our ratio of *Wolbachia* to host silent-site divergence estimates is two or three orders of magnitude lower than found for *Nasonia* or *Nomada*. This strongly supports relatively recent *Wolbachia* transfer between *D. suzukii* and *D. subpulchrella*, being inconsistent with ratios observed under cladogenic *Wolbachia* acquisition. Given that we are looking at only single *w*Spc and *w*Suz sequences, their divergence time provides an upper bound for the time of interspecific transfer (Gillespie & Langley 1979). Additional support for non-cladogenic transmission comes from the analyses of Richardson *et al.* (2012), who inferred that *Wolbachia* substitution rates were roughly ten-fold lower than the non-coding nuclear mutation rate for *D. melanogaster*, which is often considered a reasonable approximation for the rate of third-position substitutions (at least for four-fold degenerate sites, Obbard *et al.* (2012)). This is clearly inconsistent with the three-order-of-magnitude difference we estimate (Table 3).

**Table 3.** Estimated frequencies of synonymous (*k*_s_) versus non-synonymous (*k*_a_) substitutions per site for *Wolbachia* in various hosts.

| Data source | Species 1 | Species 2 | Host | | *Wolbachia* | | $k_s$ ratio |
| --- | --- | --- | --- | --- | --- | --- | --- |
| | | | $k_s$ | $k_a$ | $k_s$ | $k_a$ | |
| this work | *Drosophila suzukii* | *Drosophila subpulchrella* | $1.2 \times 10^{-1}$ | $6.8 \times 10^{-3}$ | $3 \times 10^{-5}$ | $4 \times 10^{-5}$ | 0.00025 |
| Raychoudhury et al. | *Nasonia giraulti* | *Nasonia longicornis* | $1.22 \times 10^{-2}$ | $5.4 \times 10^{-3}$ | $3.7 \times 10^{-3}$ | $2.2 \times 10^{-3}$ | 0.30 |
| Gerth & Bleidorn | *Nomada ferruginata* | *Nomada leucophthalma* | $1.95 \times 10^{-2}$ | $2.6 \times 10^{-3}$ | $2.5 \times 10^{-3}$ | $9 \times 10^{-4}$ | 0.13 |
| ibid. | *N. ferrug.* | *N. flava* | $1.92 \times 10^{-2}$ | $2.7 \times 10^{-3}$ | $2.5 \times 10^{-3}$ | $9 \times 10^{-4}$ | 0.13 |
| ibid. | *N. ferrug.* | *N. panzeri* | $1.84 \times 10^{-2}$ | $3.1 \times 10^{-3}$ | $2.7 \times 10^{-3}$ | $1.1 \times 10^{-3}$ | 0.15 |
| ibid. | *N. leuco.* | *N. flava* | $6.8 \times 10^{-3}$ | $4 \times 10^{-4}$ | $1 \times 10^{-4}$ | $1 \times 10^{-4}$ | 0.015 |
| ibid. | *N. leuco.* | *N. panzeri* | $5.8 \times 10^{-3}$ | $8 \times 10^{-4}$ | $3 \times 10^{-4}$ | $2 \times 10^{-4}$ | 0.052 |
| ibid. | *N. flava* | *N. panzeri* | $5.5 \times 10^{-3}$ | $9 \times 10^{-4}$ | $3 \times 10^{-4}$ | $3 \times 10^{-4}$ | 0.055 |

Comparing *w*Suz and *w*Spc, we found no difference in *k*_s_ and *k*_a_ (Table 1). This is also true for *w*Mel variation in *D. melanogaster* (Richardson *et al.* 2012). Gerth & Bleidorn (2016, pers. comm.) find essentially identical estimates of *k*_s_ and *k*_a_ for all pairwise comparisons of the *Wolbachia* in the clade (*(N. leucophthalma, N. flava), N. panzeri*). In contrast, comparing *w*Ri and *w*Au using the 429,765 bp dataset of single-copy, full-length genes (Table S3), we estimate *k*_s_ of 4.34%; whereas the estimated *k*_a_ is only 0.65% (or *k*_s_/*k*_a_ = 6.7). Similarly, when comparing the *Wolbachia* of the outgroup host, *N. ferruginata*, to the *Wolbachia* of the three ingroup species, Gerth & Bleidorn (2016, pers. comm.) observed *k*_s_/*k*_a_ values of 2.8, 2.8 and 2.5. In their comparisons of *w*NlonB1 and *w*NgirB from *N. longicornis* and *N. giraulti*, Raychoudhury *et al.* (2008) estimated *k*_s_/*k*_a_ = 0.0037/0.0022 = 1.7. Our data and those from other very recently diverged *Wolbachia* are consistent with either accelerated adaptive *Wolbachia* evolution in a new host or a relaxation of constraints on non-synonymous substitutions.

Estimating absolute divergence times (*i.e.*, times to the MRCA) for *w*Suz and *w*Spc and their hosts is more difficult. Assuming 10 generations per year in *Drosophila* and using the *w*Mel-derived estimate of (2.88×10^−10^, 1.29×10^−9^) changes/site/host-generation as the 95% confidence interval for the third-position substitution rate of *Wolbachia* (Richardson *et al.* 2012), *w*Suz and *w*Spc diverged about 1,600 to 7,000 years ago. Using the 95% confidence interval for first- and second-position substitution rates from Richardson *et al.* (2012) yields *w*Suz-*w*Spc divergence dates of 1,200 to 9,100 years. Given that *D. suzukii* and *D. subpulchrella* seem to be temperate species (Takamori *et al.* 2006; Ometto *et al.* 2013), the number of generations per year may be overestimated by a factor of two, which would inflate the *Wolbachia* divergence time by a factor of two. This does not affect our conclusions. Raychoudhury *et al.* (2008) estimated a *Wolbachia k*_s_ of 4.7×10^−9^ changes/synonymous site/year in *Nasonia.* Using our *k_s_* from Table 1 with the *Nasonia* calibration, the estimated divergence for *w*Suz and *w*Spc is 6,400 years, which is consistent with our *Drosophila* calibration. These analyses suggest that *w*Suz and *w*Spc diverged on the order of 1,000-10,000 years ago, orders of magnitude shorter than typical time scales for *Drosophila* speciation (10^5^-10^6^ years, Coyne & Orr 2004, p. 75; Obbard *et al.* 2012). Molecular estimates of *Drosophila* divergence times generally depend on speculative inferences from the phylogeography of the Hawaiian *Drosophila* radiation (Obbard *et al.* 2012). Using the Obbard *et al.* (2012) summary of available estimates for *D. melanogaster* and *D. simulans* divergence and our relative chronogram for *D. subpulchrella* and *D. suzukii* (Fig. 2C), we infer divergence times for *D. subpulchrella* and *D. suzukii* ranging from about one to nine million years, two orders of magnitude larger than our estimates for *w*Suz versus *w*Spc. Hence, despite great uncertainties, our data clearly preclude cladogenic transmission of *w*Suz and *w*Spc. This conclusion is further supported in the Discussion by a review of variation in rates of bacterial molecular evolution.

### Genome differences between wSpc, wSuz and wRi: structural variation and candidate genes

We identified copy-number variants (CNV) in *w*Suz and *w*Spc relative to the *w*Ri reference sequence by plotting read depth along each genome (Fig. 3; Table 4). *w*Spc and *w*Suz share a deletion relative to *w*Ri of 23,000 bp, between positions 733,000-756,000. *w*Suz has duplications 22,500 bp long from about 570,000 to 592,500 and 1,077,500 to 1,100,000. Both regions are part of the WO-B prophage. In *w*Ri, there are two nearly identical copies (99.4%) of WO-B, from about 565,000 to 636,000 and from about 1,071,000 to 1,142,000 (Klasson *et al.* 2009). *w*Suz had an additional duplication between 1,345,000 and 1,347,500, outside of the WO prophage regions (Table 4).

**FIGURE 3.**
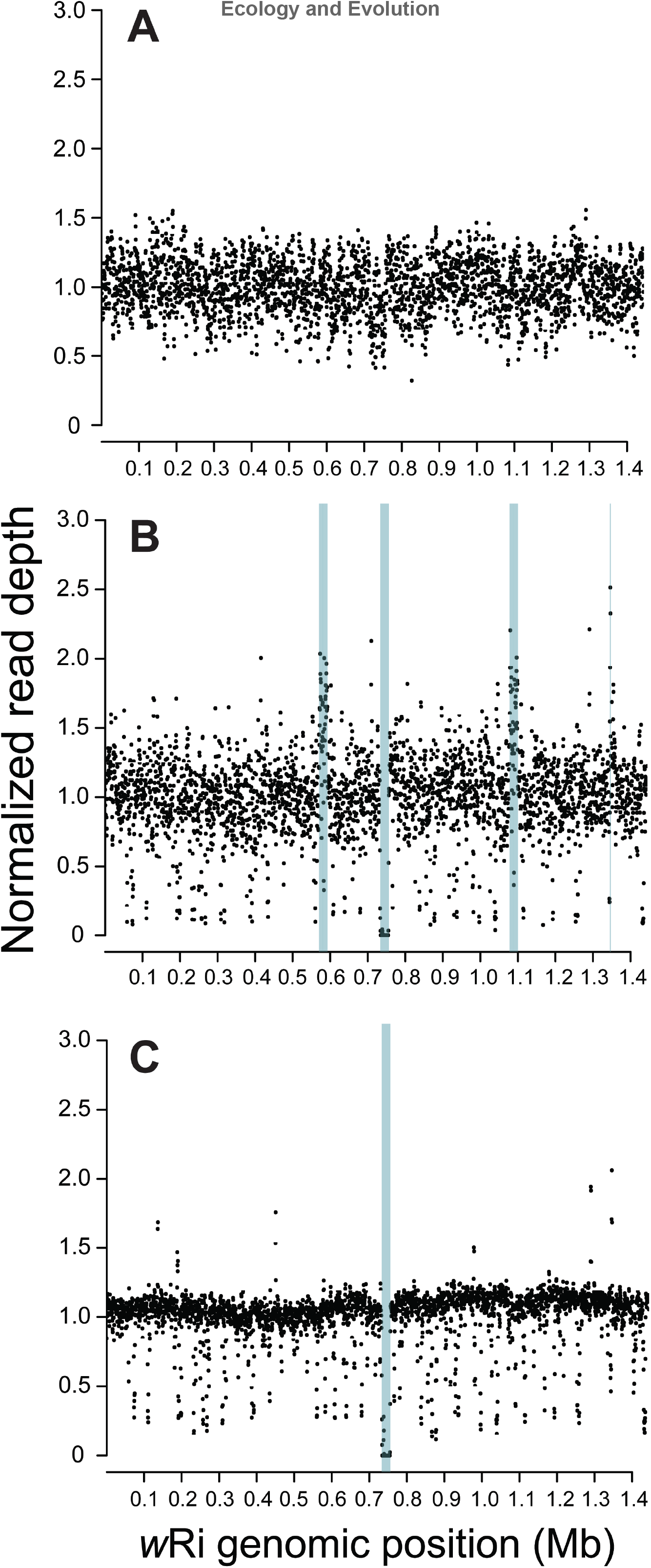
We compare normalized read-density relative to the *w*Ri reference sequence of Klasson *et al.* (2009) for: A) the Illumina reads from the Riv84 version of *w*Ri reported by Iturbe-Ormaetxe *et al.* (2010), B) the *w*Suz reads from Ometto *et al.* (2014), and C) the *w*Spc reads from this study.

**Table 4.** Copy-number variants in *w*Suz and *w*Spc relative to *w*Ri. All positions are given relative to the *w*Ri reference of Klasson *et al.* (2009).

| Start position | End position | Copy number change | Kolmogorov-Smirnov *P*-value | Affected genomes |
| --- | --- | --- | --- | --- |
| 570000 | 592500 | 2 → 3* | <0.0001 | *wSuz* |
| 733000 | 756000 | 1 → 0 | <0.0001 | *wSuz*, *wSpc* |
| 1077500 | 1100000 | 2 → 3* | <0.0001 | *wSuz* |
| 1345000 | 1347500 | 1 → 2 | 0.016 | *wSuz* |

\*This sequence is duplicated in the *wRi* genome, so it was treated as diploid in our ControlFREEC v. 8.0 analysis.

We identified homologs in our target *Wolbachia* genomes of loci implicated in producing phenotypic effects. The Octomom phenotype of *w*Mel (shortened life, high *Wolbachia* titer) has been associated with eight loci *(WD0507-WD0514*, Chrostek & Teixeira 2015, but see also Rohrscheib *et al.* 2016; Chrostek & Teixeira 2017). In the *w*Ri reference, we found homologs of only *WD0508* and *WD0509.* There were two WD0508-like genes, at 632,500-633,385 and 1,138,959-1,139,844, within the *w*Ri WO-B prophages. A single WD0509-like gene was present, from 1,419,589-1,421,396, not associated with WO-B prophage. These two genes are not neighbors in *w*Ri, *w*Spc or *w*Suz, and they are not within regions that differentiate *w*Spc and *w*Suz from *w*Ri.

Table 5 lists the orthologs and paralogs in *w*Mel, *w*Ri, *w*Suz and *w*Spc of **w*Pip_0282* and **w*Pip_0283*, the loci originally identified as CI-causing by Beckmann & Fallon (2013) in *w*Pip, the *Wolbachia* in *Culex pipiens.* These loci occur in pairs; and the “type I” pairs, orthologs of **w*Pip_0282* and **w*Pip_0283*, may be a toxin-antidote operon (compare Beckmann *et al.* 2017 with LePage *et al.* 2017). The orthologs in *w*Mel are *WD0631* and *WD0632.* As shown in Table 5, there are two copies of the type I pair in *w*Ri, one copy in each of the two complete copies of the WO-B prophage. As noted by Beckmann & Fallon (2013), in *w*Ri, there is also a paralogous pair (**w*Ri_006720* and *w*Ri_006710), termed “type II” by LePage *et al.* (2017), that exists within what they term a “WO-like island.”

**Table 5.** Homologs of CI-associated loci in *w*Mel, *w*Ri, *w*Suz and *w*Spc. The gene designations in *w*Spc and *w*Suz reflect homology to loci identified in *w*Mel and *w*Ri.

| *Wolbachia* | gene pair<sup>*</sup> | Gene 1 | Gene 2 | WO prophage association<sup>†</sup> |
| --- | --- | --- | --- | --- |
| *wMel* | I | *WD0631 (cifA/cidA)*<sup>‡</sup><br>(antidote?) | *WD0632 (cifB/cidB)*<sup>‡</sup><br>(toxin?) | yes |
| *wRi* | I.1 | *wRi_005370* | *wRi-p005380*<sup>§</sup> | yes |
|  | I.2 | *wRi_010030* | *wRi_p010040*<sup>§</sup> | yes |
|  | II | *wRi_006720* | *wRi_006710* | no |
| *wSpc* | I.1 | *wSpc_0631.I.1* | *wSpc_0632.I.1* | yes |
|  | I.2 | *wSpc_0631.I.2* | *wSpc_0632.II.2* | yes |
|  | II | *wSpc_6720* (disrupted) | *wSpc_6710* | no |
| *wSuz* | I.1 | *wSuz_0631.I.1* | *wSuz_0632.I.1* | yes |
|  | I.2 | *wSuz_0631.I.2* | *wSuz_0632.II.2* | yes |
|  | I.3 | *wSuz_0631.I.3* | *wSuz_0632.II.3* | partial<sup>¶</sup> |
|  | II | *wSuz_6720* (disrupted) | *wSuz_6710* | no |

<sup>\*</sup>Roman numerals follow the “type” designations in LePage *et al.* (2017).
<sup>†</sup>This refers to location within an intact WO prophage, as opposed to a “WO-like island” (*cf.* LePage *et al.* 2017).
<sup>‡</sup>Alternative designations (*cif* versus *cin*) from LePage *et al.* (2017) and Beckmann *et al.* (2017), respectively. Beckmann *et al.* (2017) proposes that *WD0631* produces an antidote to the toxin produced by *WD0632*.
<sup>§</sup>Annotated as pseudogenes, but see text.
<sup>¶</sup>This third copy in *wSuz* exists in the 1077500-1100000 CNV, noted in Table 4, which is a partial copy of the WO-B prophage.

Table S5 lists genes included in the CNV regions of *w*Suz and *w*Spc relative to *w*Ri. Notably, the orthologs of *WD0631* and *WD0632*, implicated in causing CI (Beckmann & Fallon 2013; LePage *et al.* 2017; Beckmann *et al.* 2017), are in a partial third copy of prophage WO-B found in *w*Suz. Hence, *w*Suz contains three copies of these two loci, whereas *w*Spc has only two (see Table 5). The copy-number variants in *w*Suz or *w*Spc do not affect the type II loci.

Table 6 reports differences among *w*Ri, *w*Suz and *w*Spc at orthologs of the CI-associated loci *WD0631, WD0632, WRi_006710*, and *WRi_006720.* The duplicate orthologs of *WD0631* in *w*Ri are *WRi_005370* and *WRi_010030.* As noted by Beckmann & Fallon (2013), the (duplicate) orthologs of *WD0632* in *w*Ri have been annotated as pseudogenes, *WRi_p005380* and *WRi_p010040*, because of premature stop codons; but they retain large, intact coding regions and may be functional (LePage *et al.* 2017 and Beckmann *et al.* 2017 provide evidence supporting function). Even with multiple orthologs of *WD0631* and *WD0632* in each genome (two in *w*Ri, two in *w*Spc, three in *w*Suz), all copies within each genome are identical and all interspecific comparisons consistently show the single-nucleotide differences reported in Table 6. *w*Suz and *w*Spc share two missense substitutions in *WD0631* and one in *WD0632.* As shown in Table 6, *w*Suz and *w*Spc also share one missense substitution in **w*Ri_006710.* This indicates that the duplications unique to *w*Suz occurred after the split of (*w*Suz, *w*Spc) from *w*Ri. *w*Spc has a nonsense mutation at position 3,353 of *WD0632*, which results in a protein lacking the last 56 amino acids produced in *w*Ri. These differences may account for the fact that while *w*Ri causes appreciable CI in *D. simulans* and detectable CI in *D. melanogaster*, neither *w*Suz nor *w*Spc causes detectable CI in its native host (Hamm *et al.* 2014).

**Table 6.** Comparisons between *w*Ri, *w*Spc and *w*Suz at the CI-associated loci (type I, possible antidote, toxin), *WD0631* and *WD0632*, from *w*Mel, and the paralogous loci (type II), *WRi_006710* and *WRi_006720* from *w*Ri. All reads from *w*Spc and *w*Suz are consistent with the differences shown.

| Location (gene, amino acid) | *wRi* codon<br>(codon, translation) | *wSpc* codon<br>(codon, translation) | *wSuz* codon<br>(codon, translation) |
| --- | --- | --- | --- |
| ***WD0631*<sup>\*</sup>** (antidote?) |  |  |  |
| 363 | AAA, Lys | GAA, Glu | GAA, Glu |
| 473 | AAA, Lys | AGA, Arg | AGA, Arg |
| ***WD0632*<sup>†</sup>** (toxin?) |  |  |  |
| 91 | GGA, Gly | GGG, Gly | GGG, Gly |
| 176 | TAT, Tyr | GAT, Asp | GAT, Asp |
| 213 | TAT, Tyr | TAC, Tyr | TAC, Tyr |
| 1118 | TTA, Leu | TGA, STOP | TTA, Leu |
| ***WRi_006710*** |  |  |  |
| 663 | TAT, Tyr | CAT, His | CAT, His |
| ***WRi_006720*** |  |  |  |
| 1 to 108 | Present | Disrupted, see text | Disrupted, see text |

<sup>\*</sup>The duplicate orthologs in *wRi* are *WRi\_005370* and *WRi\_010030*.
<sup>†</sup>The duplicate orthologs in *wRi* are *WRi\_p005380* and *WRi\_p010040*.

Our bioinformatic and PCR data show that in both *w*Spc and *w*Suz (but not *w*Ri), an IS element, identical to ISWpi7 of *w*Ri (Klasson *et al.* 2009, Table S5), has inserted before base 323 of the ortholog to *WRi_006720.* There are 21 identical copies of the ISWpi7 transposon in *w*Ri, each 1480 bp long with the transposase gene flanked on each side by about 200 bp. Clearly, this ISWpi7 insertion predates the divergence of *w*Spc and *w*Suz.

## Discussion

### Genomic data indicate non-cladogenic acquisition of wSuz and wSpc

Despite considerable uncertainly in divergence-time estimates for both *w*Suz and *w*Spc and their hosts, *D. suzukii* and *D. subpulchrella*, genomic data on relative rates of *Wolbachia* and host divergence contradict the conjecture by Hamm *et al.* (2014) that these species share similar *Wolbachia* because of cladogenic transmission. Based on this result, we must also revisit the Hamm *et al.* (2014) conclusion that cladogenic transmission of *Wolbachia* may be relatively common among *Drosophila*. That conclusion was based on the erroneous assumption that cladogenic transmission was the most plausible explanation for sister species sharing very similar *Wolbachia.* Given that on the order of half of *Drosophila* speciation events show evidence for reinforcement (*i.e*., accelerated rates of evolution for premating isolation associated with overlapping ranges) (Coyne & Orr 1989, 1997; Turelli *et al.* 2014), hybridization is apparently common among sister species of *Drosophila.* Introgression has been invoked to explain the closely related *Wolbachia* found within the *simulans* and *yakuba* clades in the *D. melanogaster* subgroup (Rousset and Solignac 1995; Lachaise *et al.* 2000). In both cases, the introgression hypothesis is favored over horizontal transmission because the hosts also share essentially identical mitochondrial DNA. *Wolbachia* transmission within the*yakuba* clade is currently being reanalyzed using complete *Wolbachia*, mitochondrial and nuclear genomes (Turelli, Conner, Turissini, Matute and Cooper, in prep.).

Understanding the frequency of alternative modes of *Wolbachia* transmission is clearly related to determining how long *Wolbachia* infections typically persist in host lineages. Bailly-Bechet *et al.* (2017) provide a meta-analysis of more than 1000 arthropod species from Tahiti that suggests average durations on the order of seven million years. However, their molecular data, which involve only two *Wolbachia* loci and the CO1 mtDNA locus, do not have sufficient power to resolve the issue. Moreover, as they note, their analysis conflates imperfect maternal transmission with the gain and loss of *Wolbachia* infections within lineages. As our analyses indicate, nearly complete *Wolbachia* and mitochondrial genomes will often be needed to unravel the acquisition and retention of closely related *Wolbachia* within host clades.

### *Extremely variable rates of* Wolbachia *molecular evolution seem an implausible alternative*

Gerth & Bleidorn (2016) proposed a time scale for *Wolbachia* evolution based on the apparent co-divergence of *Wolbachia* and nuclear genomes in a clade of four *Nomada* bee species. Our discussion of their data emphasized comparisons between the outgroup host *N. ferruginata* and the three ingroup hosts, noting that the co-divergence of these hosts and their *Wolbachia* produced relative rates of molecular divergence comparable to those inferred for a pair of *Nasonia* (Raychoudhury *et al.* 2008) and for *D. melanogaster* (Richardson *et al.* 2012). However, if we consider instead the sister species *N. leucophthalma* and *N. flava* from Gerth & Bleidorn (2016), we would infer much slower divergence of their *Wolbachia* (which recently acquired a biotin synthesis operon). For *N. leucophthalma* and *N. flava*, Gerth & Bleidorn (2016, pers. comm.) estimated synonymous nuclear substitution divergence of 6.8×10, with a corresponding *Wolbachia* synonymous substitution divergence of only 1.0×10^−4^ (Table 3). Under cladogenic transmission, this implies *Wolbachia* divergence that is roughly an order of magnitude slower than inferred from the three outgroup comparisons, with *Wolbachia* divergence at 1/68^th^ the rate of the host nuclear genomes rather than 1/8. This indicates either 8.5-fold rate variation for *Wolbachia* molecular evolution or that cladogenic transmission does not apply to this sister pair.

To explain our *D. suzukii* and *D. subpulchrella* data with cladogenic transmission and relative rate heterogeneity, we require that *Wolbachia* divergence is more than 1000-fold slower than third-position nuclear divergence. This relative rate is 100-fold slower than inferred for *D. melanogaster* and 30-fold slower than the slow rate implied by cladogenic transmission between *N. leucophthalma* and *N. flava.* Such extreme heterogeneity seems implausible, but more examples of cladogenic *Wolbachia* transmission are needed to definitively rule this out.

Although there are relatively few taxa for which we can quantify the relative rates of nuclear versus *Wolbachia* molecular evolution, there are extensive data assessing the relative constancy of bacterial molecular evolution. Kuo & Ochman (2009) provide an overview, emphasizing that variation across taxa is too great for any locus or group of loci to provide a broadly applicable “molecular clock” for bacteria. Nevertheless, their analyses indicate that variation across lineages is typically much less than 10-fold. Yet, if *w*Suz and *w*Spc were cladogenically inherited and we assume the implausibly short host divergence time of 500,000 years (half of our lowest plausible estimate, see Fig. 2), the inferred upper bound on the rate of *Wolbachia* silent site substitutions is about 1.0×10^−11^ per site per year. In contrast, the inferred rates of silent site substitutions from the *Nasonia* and *Nomada* data (Table 3) are at least two orders of magnitude faster. Such variation in *Wolbachia* substitution rates over many loci would be unprecedented among bacteria.

### Comparative genomics and cytoplasmic incompatibility

Recent experiments strongly suggest that the *w*Mel loci *WD0631* and *WD0632*, contained within the WO-B prophage, cause CI (Beckmann & Fallon 2013; LePage et al. 2017; Beckmann *et al.* 2017). Despite having orthologs of both loci that are fairly similar to those in *w*Ri, *D. suzukii* and *D. subpulchrella* show no apparent CI. There are two copies of these CI-associated loci in *w*Ri, two in *w*Spc, and three in *w*Suz. As argued above, the additional copy in *w*Suz was acquired after *w*Suz and *w*Spc diverged. The differences we document in Table 6 between *w*Ri, *w*Suz and *w*Spc at the CI-associated loci may be informative about the portions of those loci essential to CI. Unpublished data (L. Mouton, pers. comm.) show that *w*Ri causes detectable, but slight, CI when introduced into *D. suzukii.* Given the high level of CI that *w*Ri causes in *D. simulans*, these data suggest that *D. suzukii* may suppress CI, perhaps indicating a relatively old association with CI-causing *Wolbachia* (Turelli 1994; Hoffmann & Turelli 1997). We may be able to determine whether *D. suzukii* or *D. subpulchrella* was the donor of their closely related *Wolbachia* from population genomic analyses of their mtDNA and *Wolbachia*. Genomes from a geographically diverse sample of *D. suzukii* are currently being analyzed and may resolve the direction of *Wolbachia* transfer (J. C. Chiu, pers. comm.).

The published crossing studies in *D. suzukii* and *D. subpulchrella*, which found no statistically significant CI caused by *w*Suz or *w*Spc, are relatively small (Hamm *et al.* 2014; Cattel *et al.* 2016). They are comparable to the experiments that inferred no CI associated with the native *Wolbachia* infections in *D. yakuba, D. teissieri* and *D. santomea* (Charlat *et al.* 2004; Zabalou *et al.* 2004). However, larger experiments by Cooper *et al.* (2017) revealed consistent, albeit weak, CI in all three*yakuba-clade* species _ and interspecific CI between these species. More replicated assays for CI in *D. suzukii* and *D. subpulchrella*, as well as investigation of whether CI is produced when *w*Spc and *w*Suz are transinfected into CI-expressing hosts such as *D. simulans*, will indicate whether the differences described in Table 6 are candidates for disrupting the molecular processes underlying CI (Beckmann *et al.* 2017, LePage *et al.* 2017).

### Conclusions and open questions

Understanding how host species acquire *Wolbachia* requires comparing divergence-time estimates for closely related *Wolbachia* in host sister species to divergence-time estimates for both their hosts’ nuclear genes and mtDNA. To make confident inferences, we need better estimates of both the mean and variance of relative divergence rates for these three genomes. The variance for mtDNA divergence can be obtained from extant data, such as the many available *Drosophila* genomes. Estimates for nuclear, mitochondrial and *Wolbachia* genomes can be obtained from groups like the filarial nematodes for which co-divergence of the hosts and their obligate *Wolbachia* is well established (Bandi *et al.* 1998). Our ability to infer processes of *Wolbachia* acquisition will be greatly enhanced by additional examples of cladogenic transmission among insects, besides *Nasonia* wasps (Raychoudhury *et al*. 2008) and *Nomada* bees (Gerth & Bleidorn 2016). For *D. suzukii* and *D. subpulchrella*, distinguishing between introgression and horizontal transmission requires mtDNA sequences, which will be analyzed in our forthcoming *D. subpulchrella* genome paper.

It is a challenge to understand the pattern of molecular evolution between closely related *Wolbachia* whereby all three nucleotide positions evolve at similar rates, producing comparable rates of synonymous versus non-synonymous substitutions. This is consistent with the pattern of variation seen for *w*Mel within *D. melanogaster* (Richardson *et al.* 2012). In contrast, *k*_s_/*k*_a_ increases to 2-3 for the cladogenically transmitted *Wolbachia* in *Nasonia* and *Nomada*; then increases to about 7 for the more distantly related *w*Au and *w*Ri infecting *D. simulans.* Does *Wolbachia* “invasion” of a new host represent a relaxation of selective constraint or an opportunity for adaptation? The reigning paradigm for molecular evolution of endosymbionts involves the fixation of slightly deleterious mutations (Moran 1996; Kuo & Ochman 2009), consistent with relaxed constraints and reduced effective population size. However, we can test for rapid adaptation of *Wolbachia* to hosts by moving near-identical *Wolbachia* between closely related hosts and comparing fitness (and reproductive) effects in native versus novel hosts.

## Author Contributions

The genomic data for *D. subpulchrella* and *w*Spc were generated by O. R.-S., L. O., M. B. and G. A. The bioinformatic analyses were performed by W. R. C. with input from M. T., M. B. and O. R.-S. The first draft of the manuscript was produced by M. T., W. R. C. and M. B. with subsequent improvements by all authors.

## Data Accessibility

Our nuclear data from *D. subpulchrella* are available from GenBank under accession numbers MF908506-MF909523. The assembly of *w*Spc is available from GenBank under accession number XXXX.

## Acknowledgements

We thank: John Beckmann, Seth Bordenstein, Brandon Cooper, Michael Gerth, Christian Schlötterer and two anonymous reviewers for comments on previous drafts; Dave Begun, Chuck Langley, Mike May, Brian Moore and Li Zhao for help with our analyses; Shane McEvey and Antoine Abrieux for photos; and Joanna Chiu, Michael Gerth and Laurence Mouton for sharing unpublished data.

## Supporting information

Additional information may be found in the online version of this article.

**Table S1.** Near-universal, single-copy proteobacteria genes (out of 221) found using BUSCO v. 3.0.0.

| Genome | Complete, single copy | Fragment | Not found |
| --- | --- | --- | --- |
| wRi | 180 | 2 | 39 |
| wMel | 180 | 2 | 39 |
| wAu | 181 | 2 | 38 |
| wHa | 179 | 3 | 39 |
| wNo | 181 | 4 | 36 |
| wSuz | 182<sup>*</sup> | 2<sup>†</sup> | 37 |
| wSpc | 182<sup>*</sup> | 2<sup>†</sup> | 37 |

<sup>\*</sup>Ribonuclease H-like domain is complete in wSuz and wSpc, but was not found in wRi, wMel, wAu, wHa or wNo. Dihydropteroate synthase is complete in wSuz, wSpc, wAu and wNo, but was not found in wHa, wMel or wRi.
<sup>†</sup>Peptidyl-tRNA hydrolase is complete in wSuz, wSpc, and wNo, but is fragmented in wRi, wAu, wHa and wMel. Phospho-N-acetylmuramoyl-pentapeptide-transferase is fragmented in wSuz, wSpc and wHa, but is complete in wAu, wMel and wNo.

**Table S2.** Observed pairwise genomic differences between *Wolbachia* strains, given as percentage of polymorphic sites in single-copy, full-length genes present in all strains.

| wSpc-wSuz-wRi dataset (704,883 base pairs, 703 genes) |  |  |
| --- | --- | --- |
| Genomes | Number of differences | Percent difference |
| wSpc v. wSuz | 28 | 0.004% |
| wRi v. wSuz | 103 | 0.014% |
| wRi v. wSpc | 99 | 0.015% |
| wSpc-wSuz-wRi-wMel-wAu dataset (480,831 base pairs, 512 genes) |  |  |
| Genomes | Number of differences | Percent difference |
| wSpc v. wSuz | 21 | 0.005% |
| wSuz v. wRi | 62 | 0.014% |
| wSpc v. wRi | 59 | 0.014% |

**Table S3.**
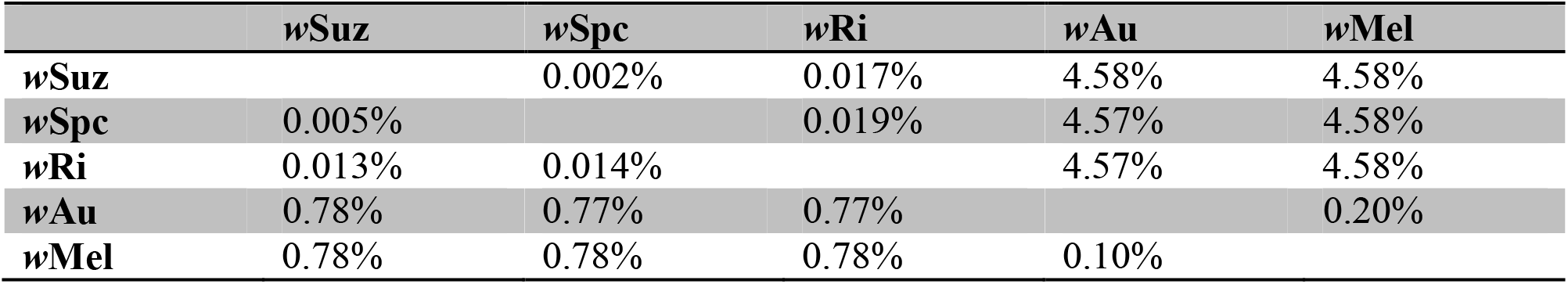
Matrix of *k*_a_ (below diagonal) and *k*_s_ (above diagonal) estimates for *w*Suz, *w*Spc, *w*Ri, *w*Au and *w*Mel (using the 480,831 bp data set from Table S2).

|  | wSuz | wSpc | wRi | wAu | wMel |
| --- | --- | --- | --- | --- | --- |
| wSuz |  | 0.002% | 0.017% | 4.58% | 4.58% |
| wSpc | 0.005% |  | 0.019% | 4.57% | 4.58% |
| wRi | 0.013% | 0.014% |  | 4.57% | 4.58% |
| wAu | 0.78% | 0.77% | 0.77% |  | 0.20% |
| wMel | 0.78% | 0.78% | 0.78% | 0.10% |  |

**Table S4.** The 28 substitutions differentiating *w*Spc and *w*Suz.

| Gene | Amino Acid | *wSpc*<br>codon | *wSuz*<br>codon | Gene Description |
| --- | --- | --- | --- | --- |
| *WRi_000230* | 228 | GTT(Val) | ATT(Ile) | DNA-directed RNA polymerase beta subunit |
| *WRi_000410* | 67 | GCT(Ala) | GCC(Ala) | hypothetical protein |
| *WRi_000410* | 118 | TCG(Ser) | TTG(Leu) | hypothetical protein |
| *WRi_000780* | 310 | TGC(Cys) | TGT(Cys) | GTP/ATP binding protein putative |
| *WRi_001670* | 103 | GAC(Asp) | GAT(Asp) | enoyl-(acyl-carrier-protein) reductase |
| *WRi_002520* | 228 | GAT(Asp) | CAT(His) | GTP-binding protein |
| *WRi_002650* | 6 | GAA(Glu) | AAA(Lys) | hypothetical protein |
| *WRi_003080* | 374 | ACA(Thr) | ACG(Thr) | succinate dehydrogenase flavoprotein subunit |
| *WRi_003240* | 59 | TCT(Ser) | CCT(Pro) | hypothetical protein |
| *WRi_003580* | 722 | TTA(Leu) | CTA(Leu) | hypothetical protein |
| *WRi_004080* | 303 | CGG(Arg) | TGG(Trp) | bicyclomycin resistance protein |
| *WRi_004790* | 793 | CCT(Pro) | TCT(Ser) | hypothetical protein |
| *WRi_004810* | 31 | AAT(Asn) | ACT(Thr) | protoheme IX farnesyltransferase |
| *WRi_006490* | 372 | AAC(Asn) | GAC(Asp) | deoxyguanosinetriphosphate<br>triphosphohydrolase |
| *WRi_006610* | 26 | CAA(Gly) | CCA(Pro) | polysaccharide deacetylase putative |
| *WRi_007380* | 335 | AAT(Asn) | AGT(Ser) | peptidase M16 family |
| *WRi_007510* | 47 | GGG(Gly) | GCG(Ala) | hypothetical protein |
| *WRi_008460* | 296 | TAT(Tyr) | CAT(His) | iron compound ABC transporter periplasmic<br>iron compound-binding protein |
| *WRi_008830* | 45 | ACT(Thr) | GCT(Ala) | hypothetical protein |
| *WRi_008830* | 29 | GCA(Ala) | GTA(Val) | hypothetical protein |
| *WRi_010700* | 179 | ATT(Ile) | ATG(Met) | permease putative |
| *WRi_010800* | 415 | TTG(Leu) | TTT(Phe) | sodium/alanine symporter family protein |
| *WRi_010800* | 226 | ATG(Met) | ATT(Ile) | sodium/alanine symporter family protein |
| *WRi_011150* | 29 | GAA(Glu) | GCA(Ala) | putative monovalent cation/H<sup>+</sup> antiporter<br>subunit D |
| *WRi_011880* | 231 | GAT(Asp) | AAT(Asn) | Succinyl-CoA synthetase beta subunit |
| *WRi_012790* | 260 | GGG(Gly) | GAG(Glu) | Type IV secretion system protein VirB9<br>putative |
| *WRi_012830* | 128 | ATT(Ile) | ATG(Met) | rod shape-determining protein RodA |
| *WRi_012980* | 22 | TGT(Cys) | TTT(Phe) | HIT family protein |

**Table S5.**
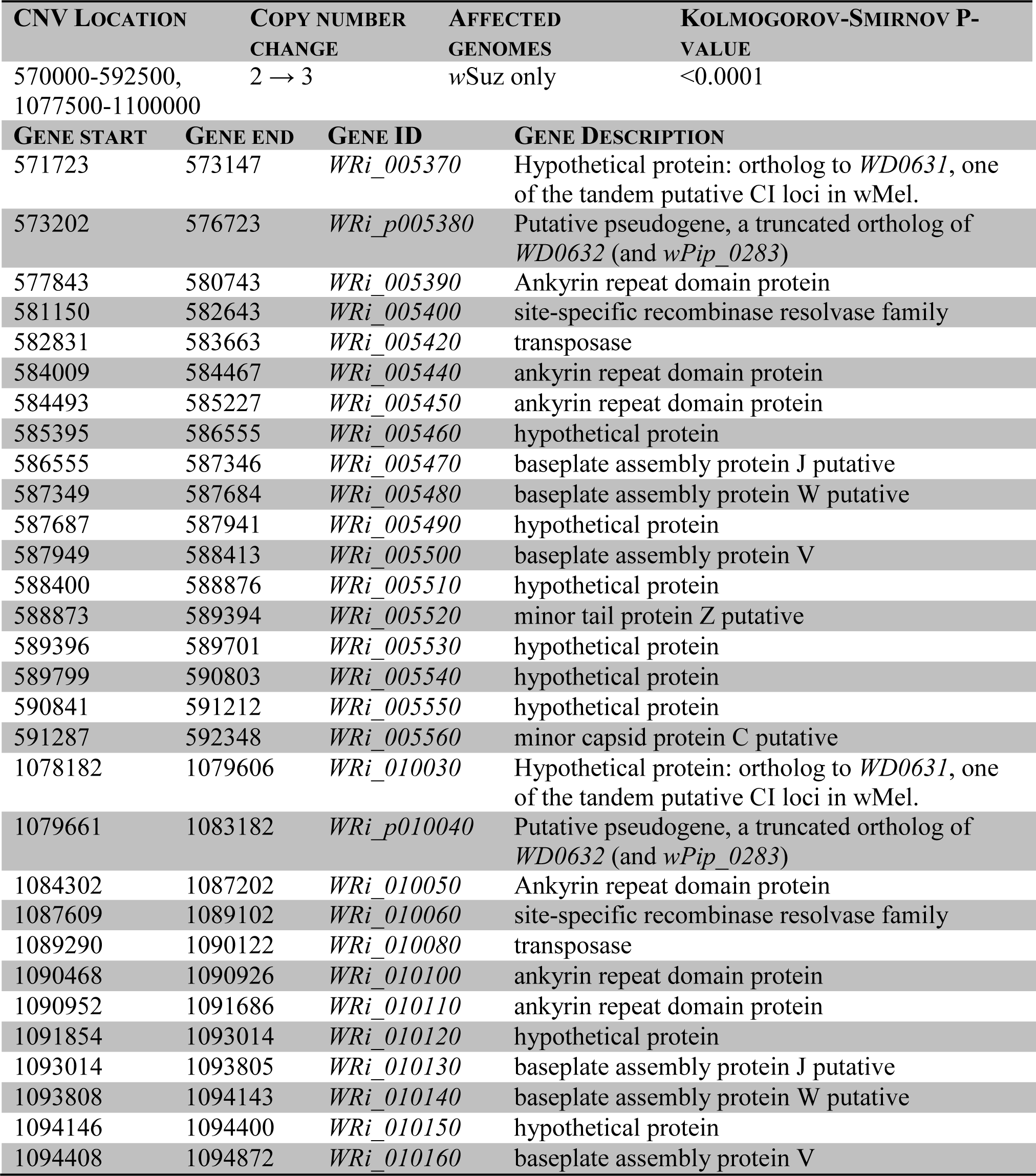
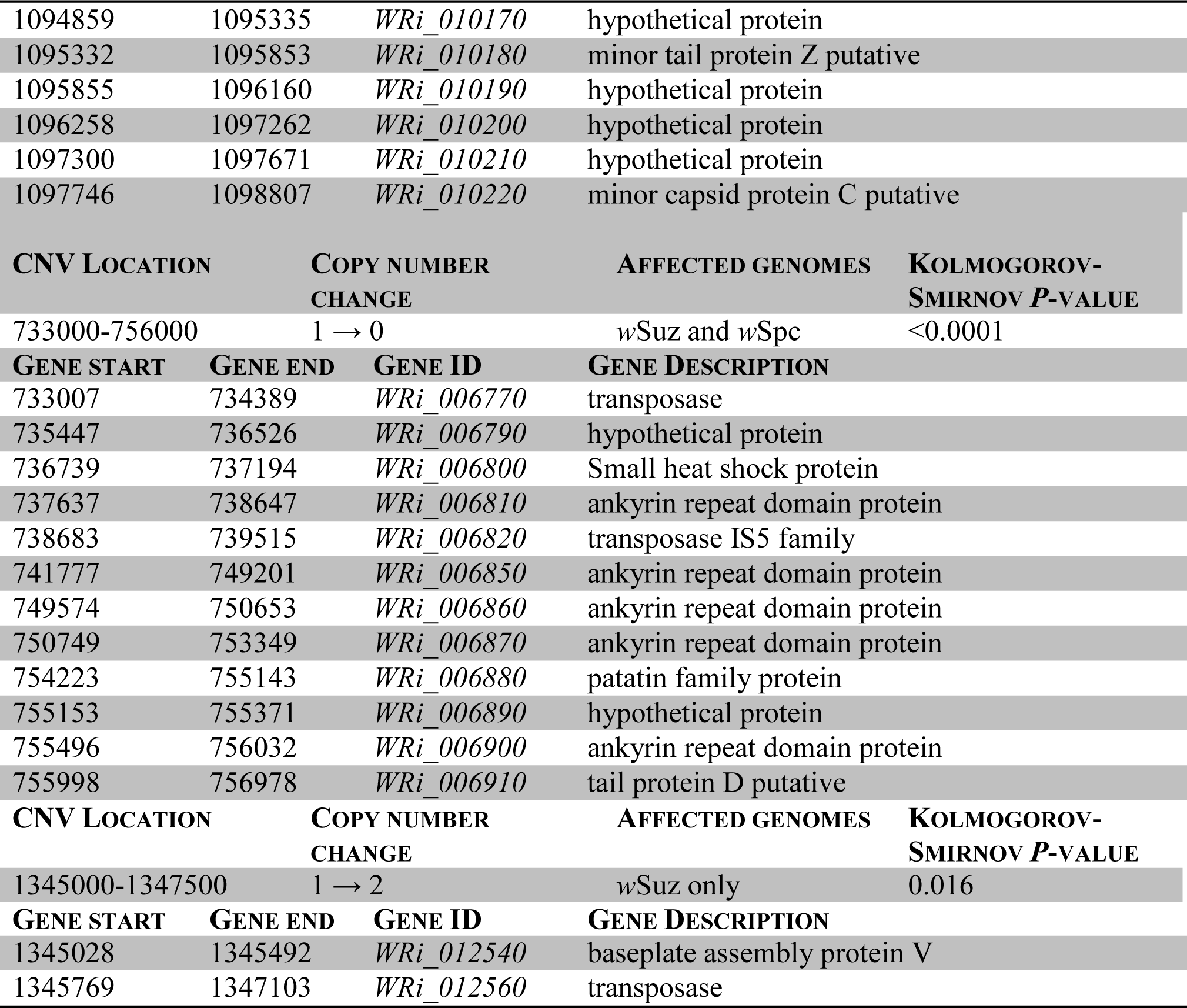
Genes present in CNV regions of *w*Suz or *w*Spc relative to *w*Ri. All locations are relative to the *w*Ri reference sequence of Klasson *et al.* (2009).

| CNV LOCATION | COPY NUMBER<br>CHANGE | AFFECTED<br>GENOMES | KOLMOGOROV-SMIRNOV P-VALUE |
| --- | --- | --- | --- |
| 570000-592500,<br>1077500-1100000 | 2 → 3 | *wSuz* only | <0.0001 |
| GENE START | GENE END | GENE ID | GENE DESCRIPTION |
| 571723 | 573147 | *WRi_005370* | Hypothetical protein: ortholog to *WD0631*, one of the tandem putative CI loci in *wMel*. |
| 573202 | 576723 | *WRi_p005380* | Putative pseudogene, a truncated ortholog of *WD0632* (and *wPip_0283*) |
| 577843 | 580743 | *WRi_005390* | Ankyrin repeat domain protein |
| 581150 | 582643 | *WRi_005400* | site-specific recombinase resolvase family |
| 582831 | 583663 | *WRi_005420* | transposase |
| 584009 | 584467 | *WRi_005440* | ankyrin repeat domain protein |
| 584493 | 585227 | *WRi_005450* | ankyrin repeat domain protein |
| 585395 | 586555 | *WRi_005460* | hypothetical protein |
| 586555 | 587346 | *WRi_005470* | baseplate assembly protein J putative |
| 587349 | 587684 | *WRi_005480* | baseplate assembly protein W putative |
| 587687 | 587941 | *WRi_005490* | hypothetical protein |
| 587949 | 588413 | *WRi_005500* | baseplate assembly protein V |
| 588400 | 588876 | *WRi_005510* | hypothetical protein |
| 588873 | 589394 | *WRi_005520* | minor tail protein Z putative |
| 589396 | 589701 | *WRi_005530* | hypothetical protein |
| 589799 | 590803 | *WRi_005540* | hypothetical protein |
| 590841 | 591212 | *WRi_005550* | hypothetical protein |
| 591287 | 592348 | *WRi_005560* | minor capsid protein C putative |
| 1078182 | 1079606 | *WRi_010030* | Hypothetical protein: ortholog to *WD0631*, one of the tandem putative CI loci in *wMel*. |
| 1079661 | 1083182 | *WRi_p010040* | Putative pseudogene, a truncated ortholog of *WD0632* (and *wPip_0283*) |
| 1084302 | 1087202 | *WRi_010050* | Ankyrin repeat domain protein |
| 1087609 | 1089102 | *WRi_010060* | site-specific recombinase resolvase family |
| 1089290 | 1090122 | *WRi_010080* | transposase |
| 1090468 | 1090926 | *WRi_010100* | ankyrin repeat domain protein |
| 1090952 | 1091686 | *WRi_010110* | ankyrin repeat domain protein |
| 1091854 | 1093014 | *WRi_010120* | hypothetical protein |
| 1093014 | 1093805 | *WRi_010130* | baseplate assembly protein J putative |
| 1093808 | 1094143 | *WRi_010140* | baseplate assembly protein W putative |
| 1094146 | 1094400 | *WRi_010150* | hypothetical protein |
| 1094408 | 1094872 | *WRi_010160* | baseplate assembly protein V |

|  |  |  |  |  |
| --- | --- | --- | --- | --- |
| 1094859 | 1095335 | WRi_010170 | hypothetical protein |  |
| 1095332 | 1095853 | WRi_010180 | minor tail protein Z putative |  |
| 1095855 | 1096160 | WRi_010190 | hypothetical protein |  |
| 1096258 | 1097262 | WRi_010200 | hypothetical protein |  |
| 1097300 | 1097671 | WRi_010210 | hypothetical protein |  |
| 1097746 | 1098807 | WRi_010220 | minor capsid protein C putative |  |
| CNV LOCATION |  | COPY NUMBER<br>CHANGE | AFFECTED GENOMES | KOLMOGOROV-SMIRNOV *P*-VALUE |
| 733000-756000 |  | 1 → 0 | wSuz and wSpc | <0.0001 |
| GENE START | GENE END | GENE ID | GENE DESCRIPTION |  |
| 733007 | 734389 | WRi_006770 | transposase |  |
| 735447 | 736526 | WRi_006790 | hypothetical protein |  |
| 736739 | 737194 | WRi_006800 | Small heat shock protein |  |
| 737637 | 738647 | WRi_006810 | ankyrin repeat domain protein |  |
| 738683 | 739515 | WRi_006820 | transposase IS5 family |  |
| 741777 | 749201 | WRi_006850 | ankyrin repeat domain protein |  |
| 749574 | 750653 | WRi_006860 | ankyrin repeat domain protein |  |
| 750749 | 753349 | WRi_006870 | ankyrin repeat domain protein |  |
| 754223 | 755143 | WRi_006880 | patatin family protein |  |
| 755153 | 755371 | WRi_006890 | hypothetical protein |  |
| 755496 | 756032 | WRi_006900 | ankyrin repeat domain protein |  |
| 755998 | 756978 | WRi_006910 | tail protein D putative |  |
| CNV LOCATION |  | COPY NUMBER<br>CHANGE | AFFECTED GENOMES | KOLMOGOROV-SMIRNOV *P*-VALUE |
| 1345000-1347500 |  | 1 → 2 | wSuz only | 0.016 |
| GENE START | GENE END | GENE ID | GENE DESCRIPTION |  |
| 1345028 | 1345492 | WRi_012540 | baseplate assembly protein V |  |
| 1345769 | 1347103 | WRi_012560 | transposase |  |

